# A multimodal imaging approach for imaging the metabolic changes resulting from bronchopulmonary dysplasia

**DOI:** 10.1101/2025.06.16.660017

**Authors:** Brittney L. Gorman, Zhi Li, Gail Deutsch, Heidi L. Huyck, Niana Beishembieva, Heather Olson, Jorge Villazon, Ping Yu, Gloria S. Pryhuber, Geremy Clair, Lingyan Shi, Christopher R. Anderton

**Author notes:** Corresponding to: Lingyan Shi, Christopher Anderton. Contributed equally to this work (Brittney L. Gorman, Zhi Li).

## Abstract

Lung tissue is composed of various functional units, each essential for maintaining the intricate functions of the lung. Disruptions in the molecular and cellular mechanisms in the lung can cause tissue fibrosis, inflammation, and severe breathing difficulties, which are common in conditions such as bronchopulmonary dysplasia (BPD). BPD’s molecular changes are not well understood, which hinders effective diagnosis and treatment. Here, we present a new multimodal imaging workflow for detailed molecular and metabolic characterization of tissues at multiple spatial scales. We applied a combined imaging approach using matrix-assisted laser desorption/ionization mass spectrometry imaging (MALDI-MSI) and ultrafast focused light-based imaging & photonics platform (U-FLIP) that included two-photon fluorescence (TPF), second harmonic generation (SHG), and stimulated Raman scattering (SRS). We also developed a hierarchical multimodal registration network (HiMReg) for the precise co-registration of each modality. This approach revealed previously unknown metabolic changes in distinct functional tissue units affected by BPD, including altered lipid distributions, reduced optical redox states, and specific collagen remodeling in bronchioles. Our findings evidenced alterations in lipid composition and metabolism of BPD-affected alveoli compared to healthy tissue, providing novel insights into disease pathophysiology. Our findings elucidate the intricate spatial and molecular complexity of BPD, building on prior research that did not provide the spatial resolution necessary to capture the nuances of metabolic alterations. This multimodal approach offers exceptional insights into disease exploration and could transform the way we study spatially heterogeneous conditions. By providing detailed maps of the metabolic shifts occurring in distinct tissue microanatomical features, the methods developed here could enable the discovery of new therapeutic avenues, making it highly attractive for the field of biomedical research.

## Introduction

Bronchopulmonary dysplasia (BPD) is a chronic lung disease that primarily affects premature infants, characterized by incomplete lung development and alveolar simplification^1^. BPD represents the predominant respiratory morbidity in preterm infants, annually affecting approximately 10,000-15,000 neonates in the United States^2,3^. Though BPD is the leading cause of morbidity among premature infants, limited treatment options exist^4^. The lung’s heterogeneity and the complex pathophysiology of BPD necessitates a comprehensive understanding of the metabolism within the lung tissue to identify potential therapeutic targets and improve patient outcomes^5^. The lung’s structural complexity provides the necessary characteristics for proper airflow and gas exchange. However, this complexity poses challenges for pinpointing molecular changes in healthy and diseased lungs that affect gas exchange when using bulk metabolomics methods^6,7^.

Traditionally, researchers have relied on biochemical assays, histological studies, and bulk tissue analyses to investigate metabolic changes, such as altered metabolism of sugars, lipids, and amino acids in BPD-affected lungs^6–10^. Recently, single-cell transcriptomics performed on BPD dissociated cells revealed the existence of an aberrant capillary cell state, uniquely found in some BPD patients^11^. While these approaches have provided valuable insights, they often lack the combination of spatial information and molecular specificity necessary to fully elucidate the intricate metabolic changes occurring across the diverse anatomical regions of the lung. Spatial proteomics using highly multiplexed immunofluorescence (MxIF) has revealed the loss of alveolar type I (AT1) cells, endothelial/capillary cells, and lymphatics, as well as an increase in alveolar type II (AT2) cells, smooth muscle, and fibroblasts^12^. While important to define changes in cell populations, MxIF, due to its targeted nature, only allows for the profiling of pre-defined protein markers and can only provide indirect and limited information on the metabolic changes occurring in the tissue. To address these limitations, several metabolic imaging technologies have emerged as powerful tools for untargeted spatial analysis of metabolites (including lipids) in biological tissues, including lung^13,14^. Moreover, the application of different metabolic imaging approaches in a multimodal fashion can overcome the limitations of using a singular imaging assay, where they can be used to provide complementary information^15^

Matrix-assisted laser desorption/ionization mass spectrometry imaging (MALDI-MSI) has been widely applied to map metabolites and identify biomarkers within tissues^16–18^. MALDI-MSI is particularly advantageous due to its ability to detect many analytes simultaneously in an untargeted fashion, while also providing precise molecular specificity^18,19^. It enables the identification of individual molecular species and offers detailed insights into molecules’ localization within distinct anatomical regions^20,21^. While MALDI-MSI can regularly achieve spatial resolutions below 10 μm, sub-cellular analyses remain challenging. Moreover, MALDI-MSI cannot provide histological information about tissues. As a consequence, MALDI-MSI is often supplemented, in a multimodal fashion, with light-based imaging methods^22–24^.

Recently, an Ultrafast Focused Light-based Imaging & Photonics platform (U-FLIP), which combines two-photon fluorescence (TPF), second harmonic generation (SHG), and stimulated Raman scattering (SRS) into one microscope, has emerged as a promising method^25–28^. This label-free optical imaging platform enables visualization of tissue morphology and can capture metabolic activity at subcellular resolutions. By detecting molecules such as NADH, FAD, and collagen, and molecular classes such as lipids and proteins, this approach can provide critical insights into metabolic and tissue morphological alterations and offers complementary information to the specific molecular annotations provided by MALDI-MSI.

Here, we introduce the integrated workflow combining U-FLIP and MALDI-MSI. We applied these methods for the comprehensive spatial analysis of distinct anatomical regions in human lung tissue affected by BPD. As part of this effort, we developed a Hierarchical Multimodal Registration network (HiMReg) to align multimodal data and permit the integrated analysis of MALDI-MSI with U-FLIP. This integration presents a unique opportunity to bridge the gap between molecular specificity and high resolution morphology. This approach evidenced functional tissue unit (FTU)-specific metabolomic changes in BPD lung tissue. By leveraging the strengths of both technologies, we characterized the lipidomic composition of alveoli, bronchioles, and vessels identified through histology staining and correlated those results with changes in lipid saturation and optical redox at high spatial resolution.

## Results

### Cross-Domain Registered Multimodal Imaging Workflow

Understanding complex and heterogenous diseases such as BPD requires comprehensive spatial characterization of tissue heterogeneity across various molecular and structural dimensions. However, integrating varying resolutions, cross-domain multimodal imaging while preserving spatial relationships has remained a major challenge in biological research. Here, we demonstrate a new multimodal imaging workflow and a novel image registration approach, enabling multi-technique correlations between molecular distributions, metabolic states, and tissue architecture within the same pulmonary tissue sections.

Our method combines MALDI-MSI for spatial lipidomics and ultrafast focused light-based imaging for metabolic imaging and histological assessment (**Figure 1**). This multi-scale analysis workflow helped to decipher the complex pathophysiology of the lung, where cellular metabolism, lipid composition, and tissue architecture are intricately correlated. A technical barrier in developing this workflow was cross-modality substrate compatibility. While conventional MALDI-MSI applications typically use electrically conductive indium tin oxide (ITO)-coated slides for optimal sensitivity, these coated substrates generated interfering nonlinear optical effects that compromised the transmission-based laser setup for SRS measurements **(Figure S1)**.

**Figure 1.**
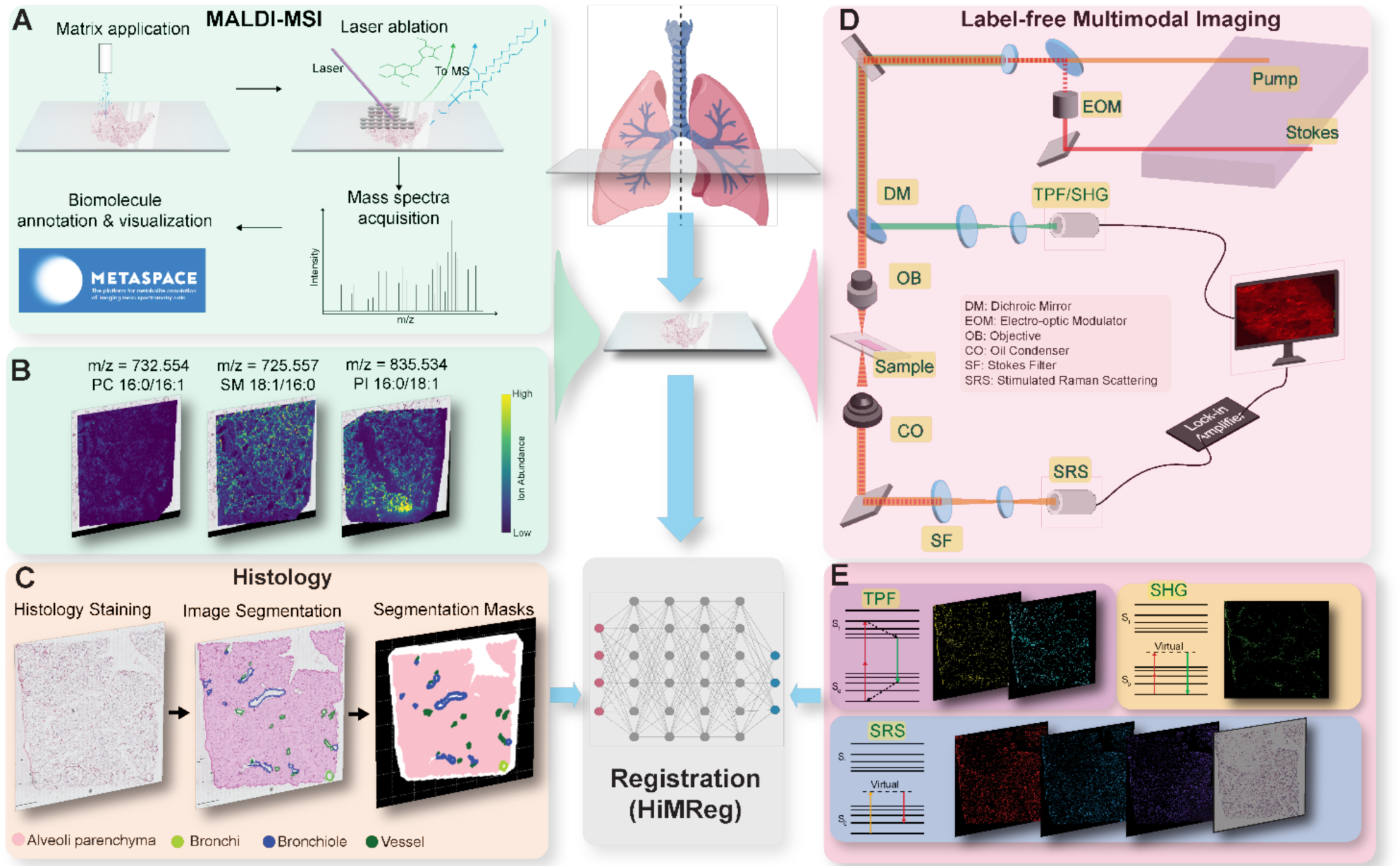
A co-registered multimodal imaging workflow for comprehensive human lung tissue analysis. (A) MALDI-MSI approach, including matrix application via automated sprayer, laser ablation for mass analysis of generated ions, and data processing using METASPACE for biomolecule annotation and visualization. (B) Representative MALDI-MS images showing spatial distribution of specific lipids in human lung: phosphatidylcholine (PC 16:1/16:0), sphingomyelin (SM 16:1/16:0), and glucosylceramide PI 16:0/18:1. (C) Workflow for histological analysis. H&E staining is performed on the same tissue section after label-free multimodal imaging. Image segmentation identifies key functional tissue units: alveoli parenchyma (pink), bronchi (green), bronchioles (blue), and vessels (dark green). A registration network (HiMReg) integrates cross modality images for comprehensive spatial analysis. (D) Schematic of U-FLIP imaging platform combining TPF, second SHG, and SRS in one microscope platform. (E) Representative images obtained from each modality in (C), where TPF captures NADH and FAD autofluorescence, SHG visualizes collagen fibers, and SRS provides chemical bonding information specific to proteins and lipids.

We found that analyzing serial sections, where consecutive tissue sections were placed on ITO-coated and normal glass slides, was sufficient for individual quantitative analyses. However, this approach was inadequate for precise spatial correlation across modalities, as it is extremely difficult to obtain two serial sections without substantial morphological and cell type changes, especially when the sections are generated from pulmonary tissues with a large proportion of open, air-filled space. Therefore, it is ideal to perform multimodal imaging on the same section to enable full integrability of datasets. We optimized MALDI-MSI parameters for non-conductive glass slides to prevent signal suppression caused by charge accumulation. The use of glass slides allowed true same-section multimodal imaging, where MALDI-MSI was performed on the tissue section first, followed by U-FLIP and histology staining.

We applied MALDI-MSI (**Figure 1A**) and annotated the resulting molecular profiles using METASPACE against the Swisslipids and our internal lung LC-MS/MS databases created using 200 µm tissue sections from the same tissue blocks. This revealed distinct lipid distributions within the lung tissue (**Figure 1B**). Post-MALDI-MSI, the matrix was removed, and the tissue sections were chemically fixed using paraformaldehyde. The same tissue section was then imaged by U-FLIP, which combines TPF for metabolic imaging, SHG for collagen visualization, and SRS for chemical composition analysis (**Figure 1C, D)**. Finally, hematoxylin and eosin (H&E) staining enabled detailed histological assessment and annotation of key FTUs for downstream quantitative analyses (**Figure 1E**).

However, achieving precise spatial registration between these diverse imaging modalities presented a significant challenge, particularly due to image distortion and disparate spatial resolutions. To address this, we developed HiMReg, which is an innovative registration framework to co-register all our imaging modalities’ data **(Figure S2A)**. This approach uses H&E-stained images as a common reference point (i.e., the anchor image that we registered with all other data). While MALDI-MSI data are regularly aligned with H&E^23,29^, the registration between high-resolution U-FLIP and H&E images posed more substantial technical hurdles. HiMReg first employed hierarchical affine registration to establish global positioning, followed by hierarchical diffeomorphic registration to account for local, non-rigid tissue deformations. The network’s multi-scale processing approach, implemented through PyTorch-based GPU acceleration^30^, minimized local minima effects while efficiently handling large-scale image processing. Alignment evaluation demonstrated HiMReg significantly outperforms traditional registration methods like Elastix^31^ **(Figure S2B,C)**, achieving superior alignment accuracy as measured by both mutual information^32^ and dice coefficient metrics^33^ **(Figure S2D,E)**.

### Spatial Lipidomics of Lung Tissue

MALDI-MSI was employed on each lung tissue section to obtain lipid composition in an untargeted fashion. Multiple tissue blocks (n=4) were selected from the left upper lobe of a healthy lung and spatially matched with blocks similar to those of a BPD-affected lung (**Table S1**). Tissue sections were first assessed using autofluorescence microscopy to locate regions of interest (ROIs) containing the four FTUs, parenchyma, bronchioles, bronchi, and large vessels (i.e., vasculature excluding the capillaries located in the alveolar parenchyma) (**Figure S3**). For each tissue, ROIs were selected containing at least one bronchi or bronchiole, vessel, and alveoli-containing region. MALDI-MSI data were acquired within each ROI at 35 µm lateral resolution to permit the identification of distinct lipid distributions within each airway, vessel, and alveolar region (**Figure 2C**).

**Figure 2.**
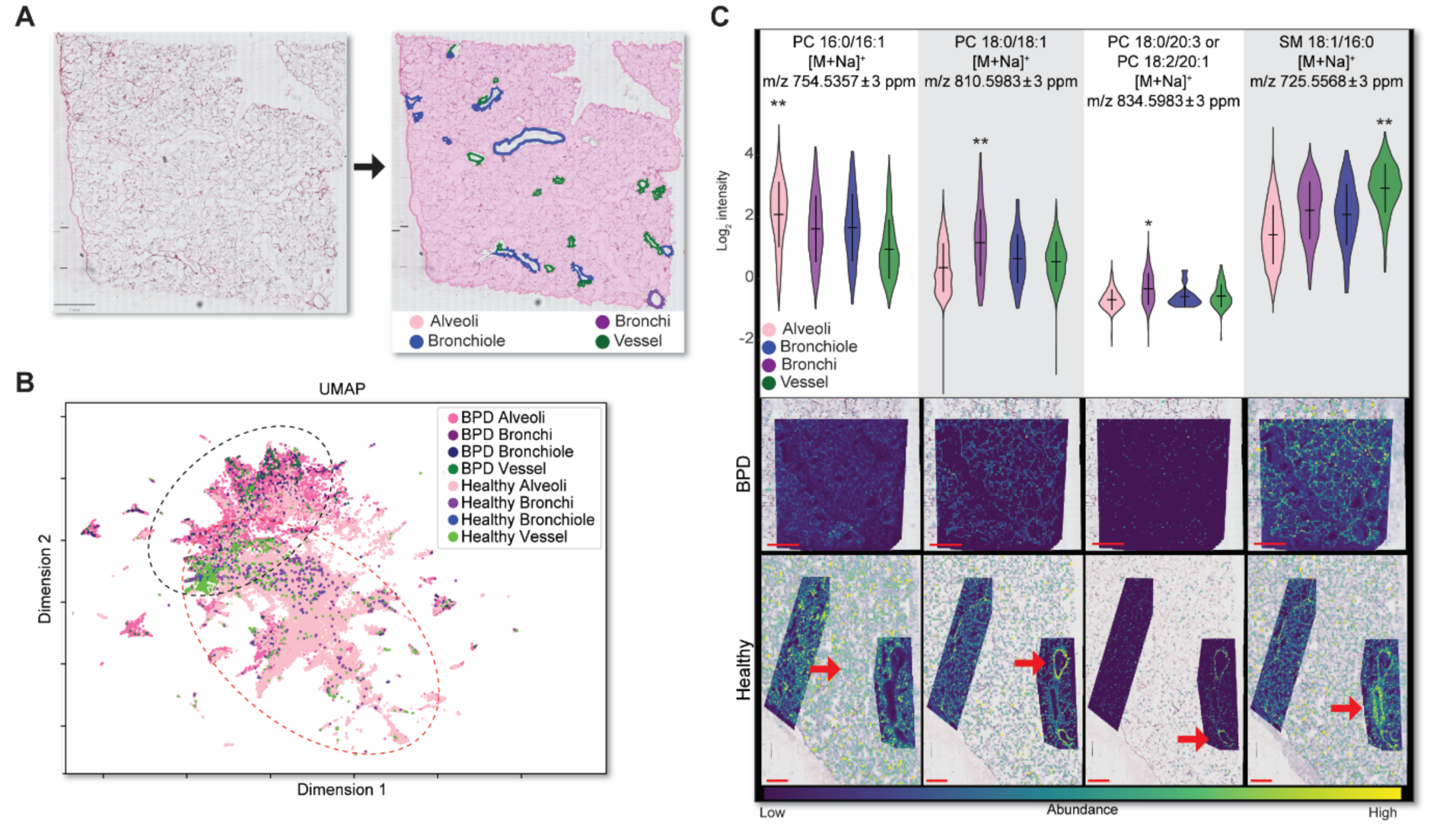
Multivariant analysis of spatial lipidomics of Healthy and BPD lung tissue. (A) H & E stained tissue section and QuPath segmentation of the alveoli, bronchi, bronchioles, and vessels. (B) UMAP analysis of the lipid annotations shows separation between Healthy and BPD lung FTUs. The most distinct separation in this multivariant analysis was between the alveoli. (C) Violin plots and corresponding ion images showing the distribution of lipids within distinct FTUs of healthy and BPD lung tissue. Red arrows indicate the position of the corresponding FTU where each ion is statistical elevated (*p<0.05, **p<0.001). Scale bar: 1 mm.

Leveraging co-registered histology, we created histology-based assessments of the lipid populations in each FTU of interest. First, we performed a multivariant analysis using a uniform manifold approximation and projection (UMAP), shown in **Figure 2C**. By graphing each analyzed pixel in the datasets according to UMAP dimensions 1 and 2, we observed modest clustering of each FTU. Notably, the alveoli-containing pixels in the healthy and BPD samples separated most significantly on the UMAP, indicating the presence of more significant disease-dependent lipidomic differences in the alveoli. To elucidate the lipids influencing the differences detected in multivariant analysis, we performed Student’s t-tests and compared the ions within each segmented FTU to all other regions in the lung. Representative lipids are shown in **Figure 2C**. Phosphatidylcholine (PC) 16:0/16:0, was most abundant in the alveoli, whereas PC 18:0/18:1 is more abundant in the bronchi and bronchioles, and sphingomyelin (SM) 18:1/16:0 was mostly present in large vessels.

Further exploration of the lipidomic signatures was done at increasing levels of chemical specificity for each FTU and comparing the healthy FTUs to the BPD ones. **Figure 3A** shows the relative abundances of the lipids organized by their subtype in each FTU. Here, we observed increased abundances of various phosphatidylcholines (PCs) in the alveoli and bronchi of the healthy tissue. Meanwhile, the annotated blood vessels displayed lower abundances of several PCs and increased abundances of sphingolipids (SLs). In particular, as shown in **Figure 3B**, we observed that PC 16:1/16:0, PC 14:0/16:0, and PC 16:0/16:0 were most abundant in the alveoli (blue arrow). The bronchi and bronchioles displayed higher abundances of PC 18:0/18:1 and PC 16:0/18:1 (green arrow). The sphingomyelin, SM 18:1/16:0, was the most intense ion identified within the vessels (red arrow). Representative ion images for each region are shown in **Figure 3C**.

**Figure 3.**
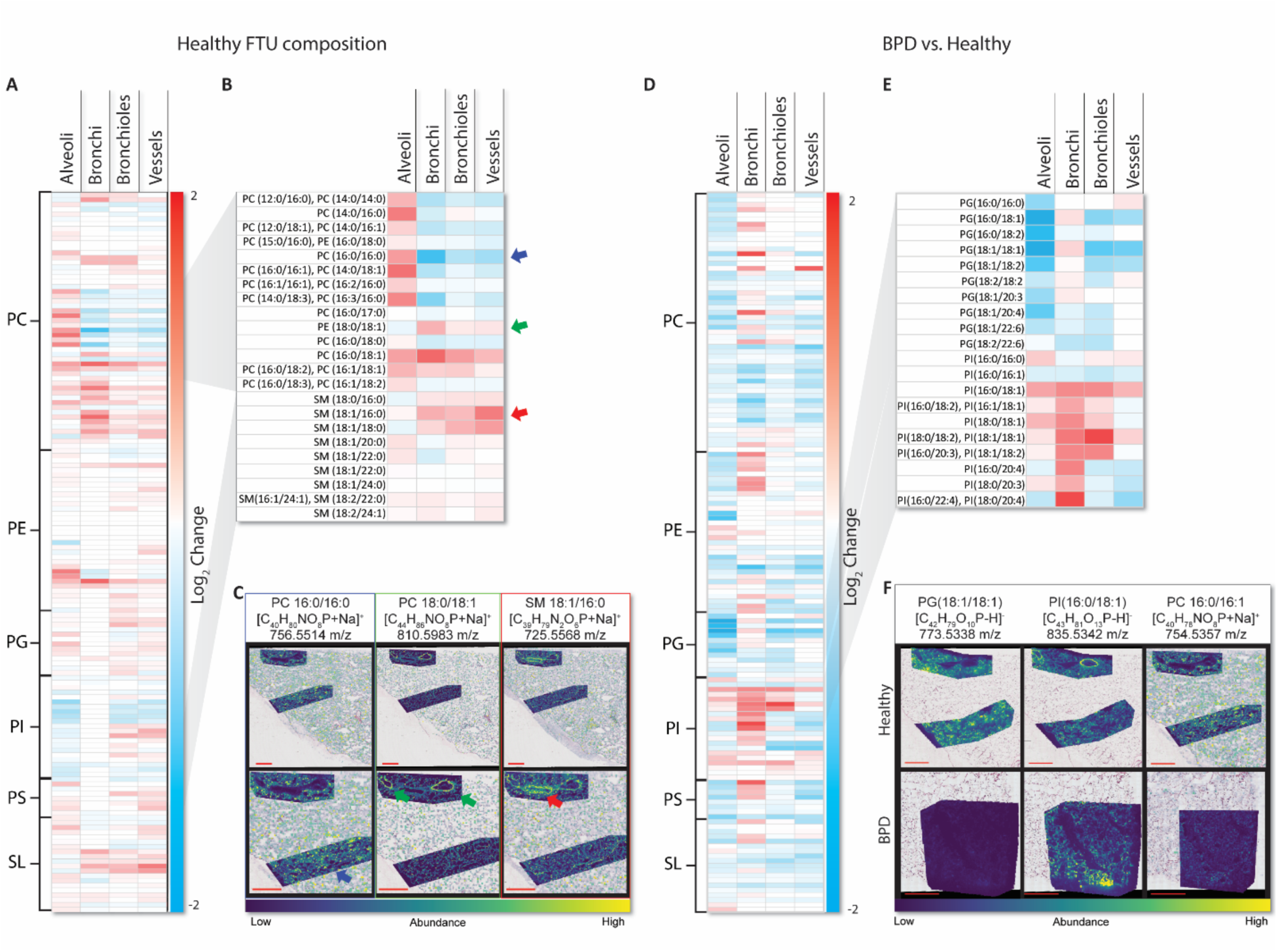
MALDI-MSI lipid profiles of healthy and BPD anatomical regions. (A) Heat map showing the relative abundance of select lipids within each segmented Healthy FTU organized by lipid subtype and increasing fatty-acid chain length from top to bottom. (B) Specific lipid annotation information for a selection of PC and SM lipids. (C) MALDI-MSI images showing the relative abundances of PC 16:0/16:0, PC 18:0/18:1, and SM18:1/16:0. PC 16:0/16:0 are the most abundant in the alveoli (blue arrow), PC 18:0/18:1 increased in the bronchi (green arrow), and SM 18:1/16:0 displayed highest association with the vasculature (red arrow). (D) Heat map showing the relative change in abundance between healthy and BPD tissues within each segmented FTU organized by lipid subtype and increasing fatty-acid chain length from top to bottom. (E) Specific lipid annotation information for a selection of PG and PI lipids. (F) MALDI-MSI images show the relative abundances of PG 18:1/18:1, PI 16:0/18:1, and PC 16:0/16:1. Scale bars are 2 mm.

When compared to the healthy tissue the BPD lung tissue displayed an overall decrease in PCs, particularly in the alveoli (**Figure 3D, E**). We also observed a decrease in phosphatidylglycerols (PGs). The bronchi and bronchioles of the BPD lung tissue displayed the highest abundances of PCs with longer fatty acid chains. In the airways of the BPD tissue, we observed an increase in phosphatidylinositol (PI) and phosphatidylethanolamine (PE) lipids. These lipidomic differences suggest complex metabolic changes in both alveolar and airways regions, with more pronounced changes in the alveolar ones. However, our MALDI-MSI analysis did not resolve other metabolic changes.

### Metabolic and Morphological Imaging of Lung Tissue

To complement the spatial lipidomic findings described above, we employed U-FLIP, enabling the visualization and quantification of key metabolic indicators and structural components^34,35^ across FTUs in healthy and BPD-affected lung tissues at high spatial resolution.

Our U-FLIP focused on several critical endogenous molecules involved in metabolism: FAD, NADH, saturated fatty acids (SFAs), and unsaturated fatty acids (USFAs); as well as protein content and morphological profiles of collagen fibers (**Figure 4A**). The normalized optical redox ratio, defined herein as FAD/(NADH + FAD), was obtained from TPF images of said molecules^36^, and lipid unsaturation ratio, defined as USFAs /(SFAs + USFAs), was derived from SRS ratiometric images of SFAs and USFAs at 2,880 and 3,011 cm^-1^, respectively^-^^37^. The optical redox ratio can be used as an indicator of cellular metabolic state, and may reflect primarily mitochondrial metabolism as the TCA cycle is the major contributor to cellular NAD+/NADH and FAD/FADH2 pools^38^. Therefore, changes in this ratio potentially offer insights into alterations in mitochondrial function and cellular energy metabolism in diseased states^39^. We investigated each spatial metabolic indicator in the segmented FTUs of healthy human lung tissue (**Figure 4B, C**), and detected significant variations across FTUs through quantitative analysis of the optical redox ratio (**Figure 4D**). In the control tissue, the annotated vessels exhibited a higher optical redox ratio than the alveoli and bronchioles, suggesting higher energy demand and oxidation in these regions. Another explanation is lower FAD(H)/NAD(H) pools persist in vascular structures. Concurrently, alveolar and vascular regions displayed a relatively higher degree of unsaturation compared to bronchioles (**Figure 4E**).

**Figure 4.**
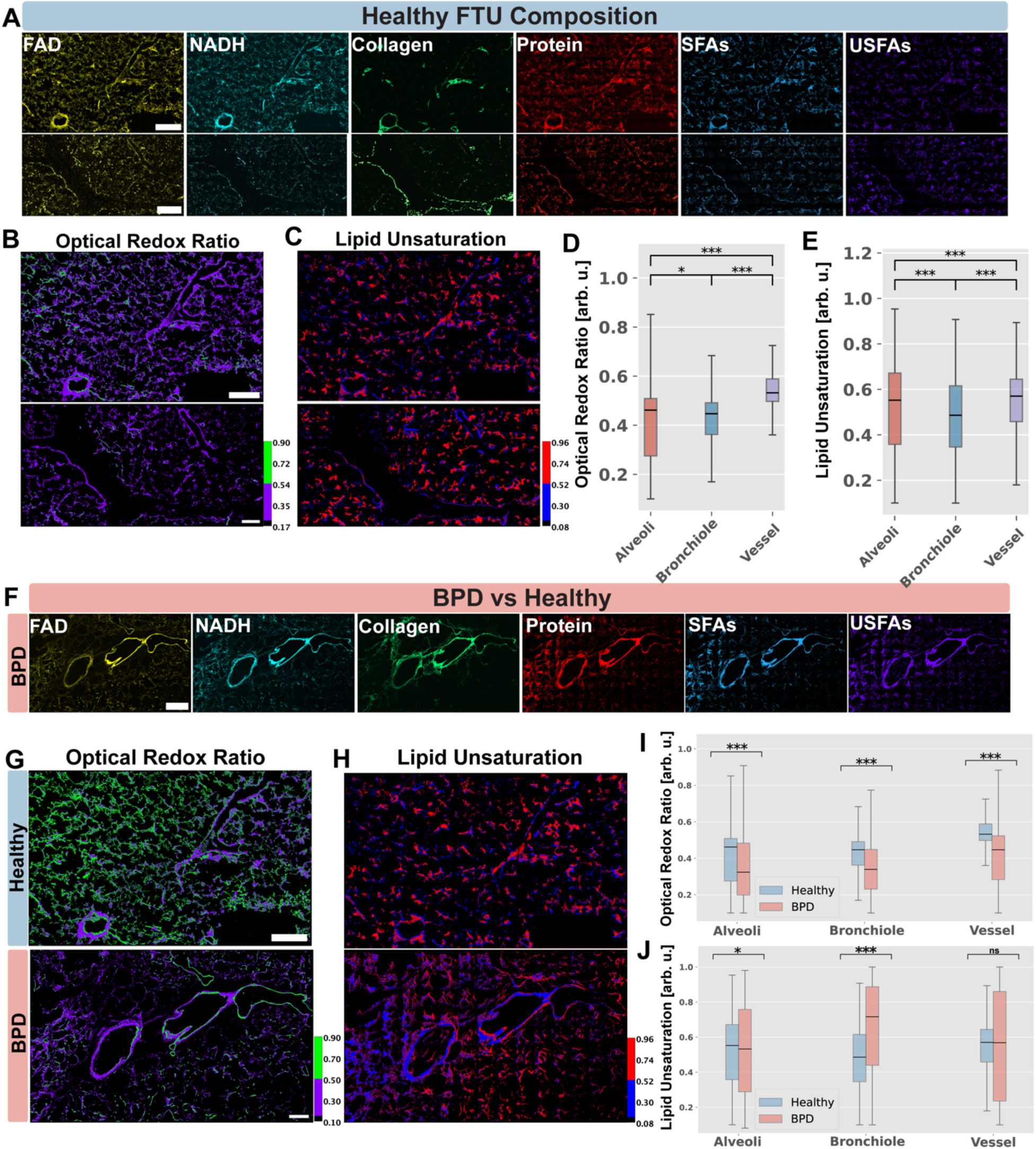
Label-free U-FLIP reveals metabolic and structural differences between healthy and BPD lung tissues. **(A)** Label-free imaging of key biomarkers in healthy FTUs. **(B-C)** Optical redox ratio and lipid unsaturation maps in healthy tissue. **(D-E)** Quantification of redox ratio and lipid unsaturation across healthy FTUs. **(F)** Comparison of biomolecular distributions in BPD vs healthy tissue. **(G-H)** Optical redox ratio and lipid unsaturation maps comparing BPD and healthy tissues. **(I-J)** Quantitative comparison of redox ratio and lipid unsaturation between BPD and healthy FTUs. Scale bar: μm. *p<0.05, ***p<0.001, ns: not significant.

Analysis of healthy versus BPD tissues revealed significant metabolic differences related to disease (**Figures 4F-J**). The optical redox ratio analysis (**Figure 4G, I)** displayed a significant decrease in the BPD tissue across all FTUs, suggesting either an alteration of the redox state or an overall reduction of the energy metabolism of the cells. Lipid unsaturation analysis comparison between healthy and BPD (**Figure 4H, J)** was more intricate. While lipid unsaturation in the alveoli was decreased compared to healthy tissue, it was increased in bronchioles. Metabolic changes in the lung remained difficult to fully decipher at the resolution of SRS.

To characterize these metabolic alterations with higher spatial granularity, we employed hyperspectral SRS (hSRS) with computational approaches to analyze the molecular composition of alveoli at subcellular resolution. Unsupervised UMAP clustering followed by k-means analysis of the SRS data revealed six distinct chemical clusters with differential enrichment between healthy and BPD (**Figure 5A, B)**. Healthy alveoli showed predominant enrichment in clusters 2 (green) and 5 (magenta), characterized by spectra indicative of obvious unsaturated lipid content while BPD alveoli exhibited higher proportions of clusters 1 (red), 3 (blue), and 4 (yellow). Cluster-specific Raman spectra demonstrated reduced unsaturated fatty acid content in BPD-enriched clusters (**Figure 5C**). Subcellular resolution chemical mapping revealed substantial differences in the molecular spatial distribution between healthy and BPD alveoli. Furthermore, using PRM-SRS^40^, a label-free lipid subtype detection method, we also observed decreased phosphatidylcholine (PC) levels and increased triglyceride (TG) accumulation in BPD alveoli (**Figure 5D).** Detailed examination of alveolar architecture revealed that cluster 4, characterized by its more saturated lipid spectral profile, primarily localized to the outer layers in both conditions but exhibited a greater thickness in BPD. While cluster 6 (cyan) persisted in the inner layers of both healthy and BPD alveoli, the inner regions of healthy alveoli were enriched in clusters 2 and 5, whereas BPD tissue showed predominant localization of clusters 1 and 3 (**Figure 5E**), indicating a potentially multicellular metabolic remodeling during disease progression.

**Figure 5.**
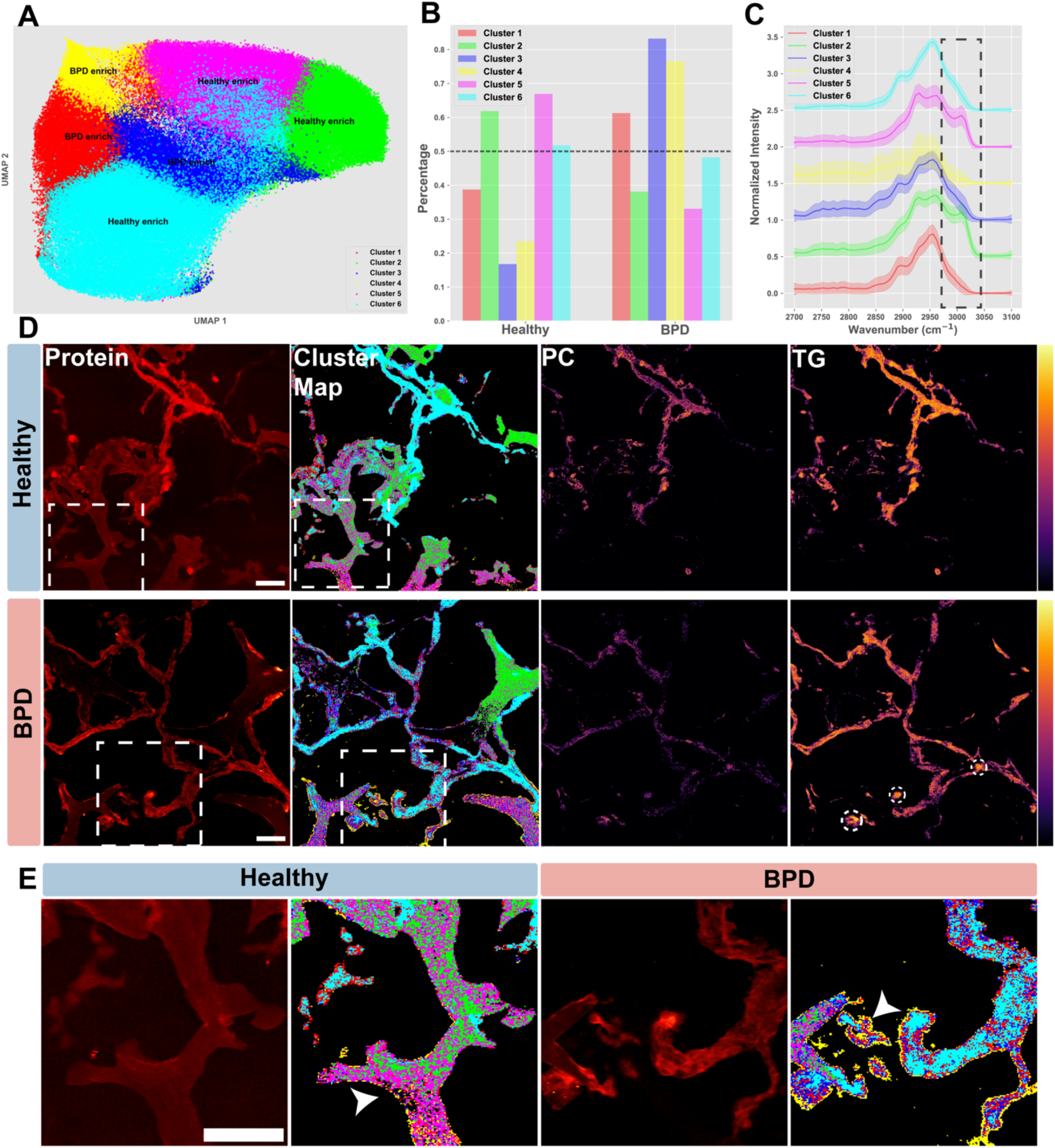
Spatial chemical profiling reveals distinct molecular compositions in healthy and BPD alveoli through hyperspectral SRS (hSRS). **(A)** Combined UMAP visualization and k-means clustering (k=6) of hSRS data, where each point represents a Raman spectrum in CH region from a tissue pixel, showing chemically distinct regions enriched in either healthy or BPD. **(B)** Percentage distribution of spectral clusters between healthy and BPD. **(C)** Normalized Raman spectra (2700-3100 cm⁻¹) for each cluster showing distinct chemical compositions. **(D)** Comparison of protein distribution (2930 cm⁻¹), cluster maps, phosphatidylcholine (PC) and triglyceride (TG) content between healthy and BPD. **(E)** High magnification views from the white dash box in (D) showing detailed molecular changes in healthy versus BPD. Scale bars are 50 μm.

Next, we investigated the structural consequences of BPD on the extracellular matrix. Collagen fibers play a crucial role in maintaining the structural integrity and mechanical properties of lung tissue. They are particularly sensitive to changes in cellular metabolism and oxidative stress^41^. Using SHG microscopy to visualize collagen architecture, we observed that the BPD lung tissue exhibited changes in collagen ultrastructure (**Figure 6A, B)**. Detailed investigation (**Figure 6C, D)** revealed distinct patterns of collagen reorganization in the different FTUs of the BPD lung. In healthy tissue, collagen appeared densely packed with well-organized fibrillar structures and concentrated distribution patterns. In contrast, BPD tissue showed more diffuse collagen arrangements, suggesting altered matrix organization. Quantitative analysis demonstrated a significant reduction in organized bronchiole-associated collagen fiber density as measured by SHG microscopy in BPD compared to healthy tissue (**Figure 6E**). This finding likely reflected alterations in collagen organization rather than a decrease in total collagen content, as previous studies showed increased collagen subunit abundance in these regions^42^. The reduced SHG signal suggested a shift from well-organized collagen fiber structures to more disorganized collagen deposition patterns in BPD tissue. Interestingly, vascular-associated collagen displayed remarkably different patterns. Despite the pronounced changes in bronchiole regions, both the density and thickness of vessel-associated collagen remained unchanged between healthy and BPD tissues (**Figure 6E, F)**.

**Figure 6.**
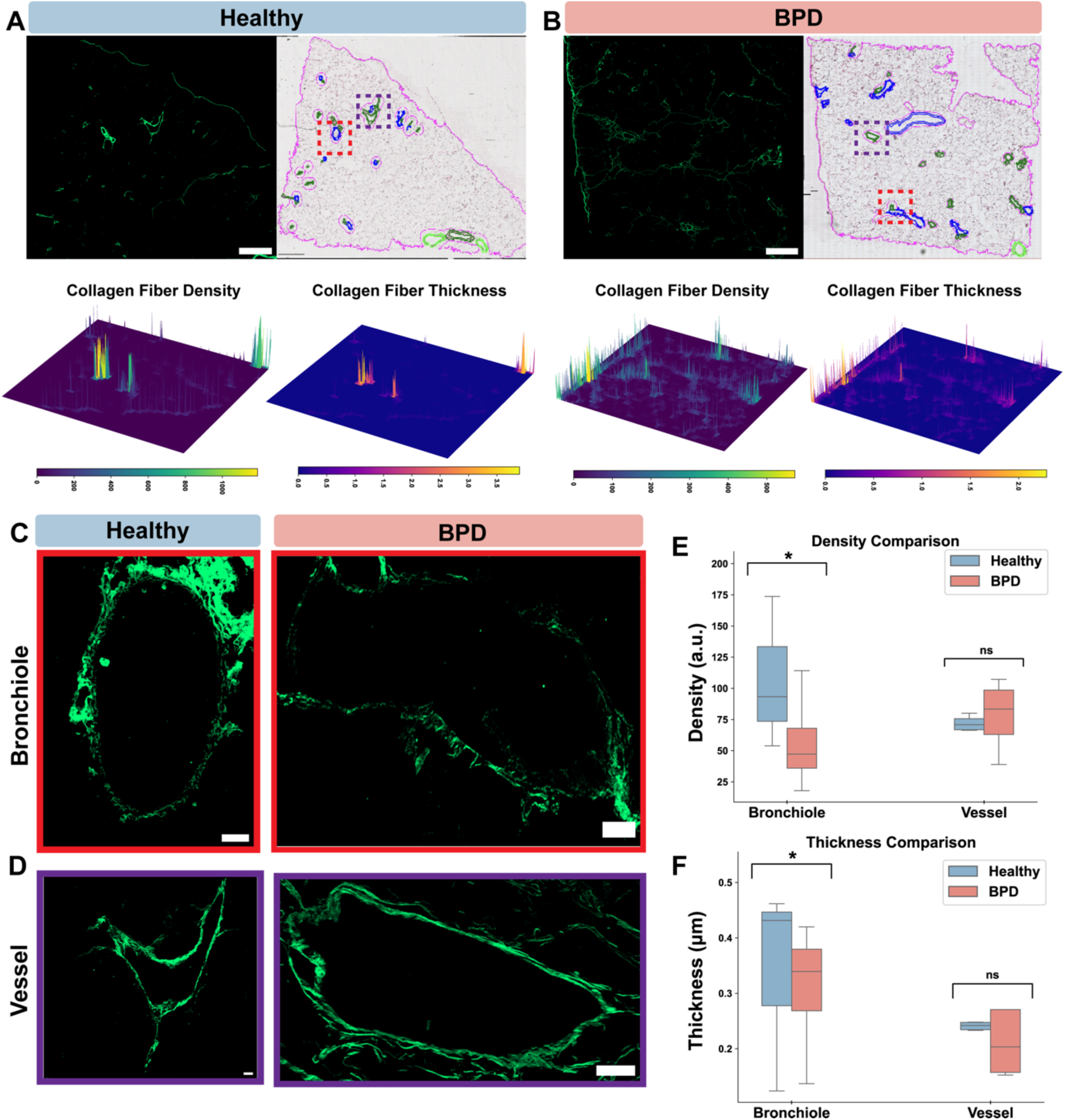
Collagen fiber analysis in healthy and BPD-affected lung tissues. (A, B) Representative SHG images and corresponding H&E stains of healthy (A) and BPD (B) lung tissues. Scale bars are 1 mm. Collagen fiber density and thickness heat maps are shown below. (C) High-magnification SHG images of bronchioles in healthy (left) and BPD (right) tissues. Scale bars: 100 μm. (D) High-magnification SHG images of vessels in healthy (left) and BPD (right) tissues. Scale bars: 100 μm. (E) Quantitative comparison of collagen fiber density in bronchioles and vessels between healthy and BPD tissues. (F) Quantitative comparison of collagen fiber thickness in bronchioles and vessels between healthy and BPD tissues. Data are presented as mean ± SD. *p < 0.05, ns: not significant.

### Co-registered MALDI-MSI and U-FLIP Correlation

Our image registration method developed and employed here has multiple significant benefits. First, it permits alignment of all imaging modalities with histology images and image segmentation. Second, the simultaneous observation of high-resolution U-FLIP images and high chemical specificity MALDI-MS images. For example, we observed collagen-rich airways associated with specific lipids (i.e., SM 18:1/18:0) (**Figure 7A**). Finally, it enhances our targeted analyses of the lipidome by providing regions of interest based on specific metabolic readouts, such as lipid unsaturation or optical redox ratio.

**Figure 7.**
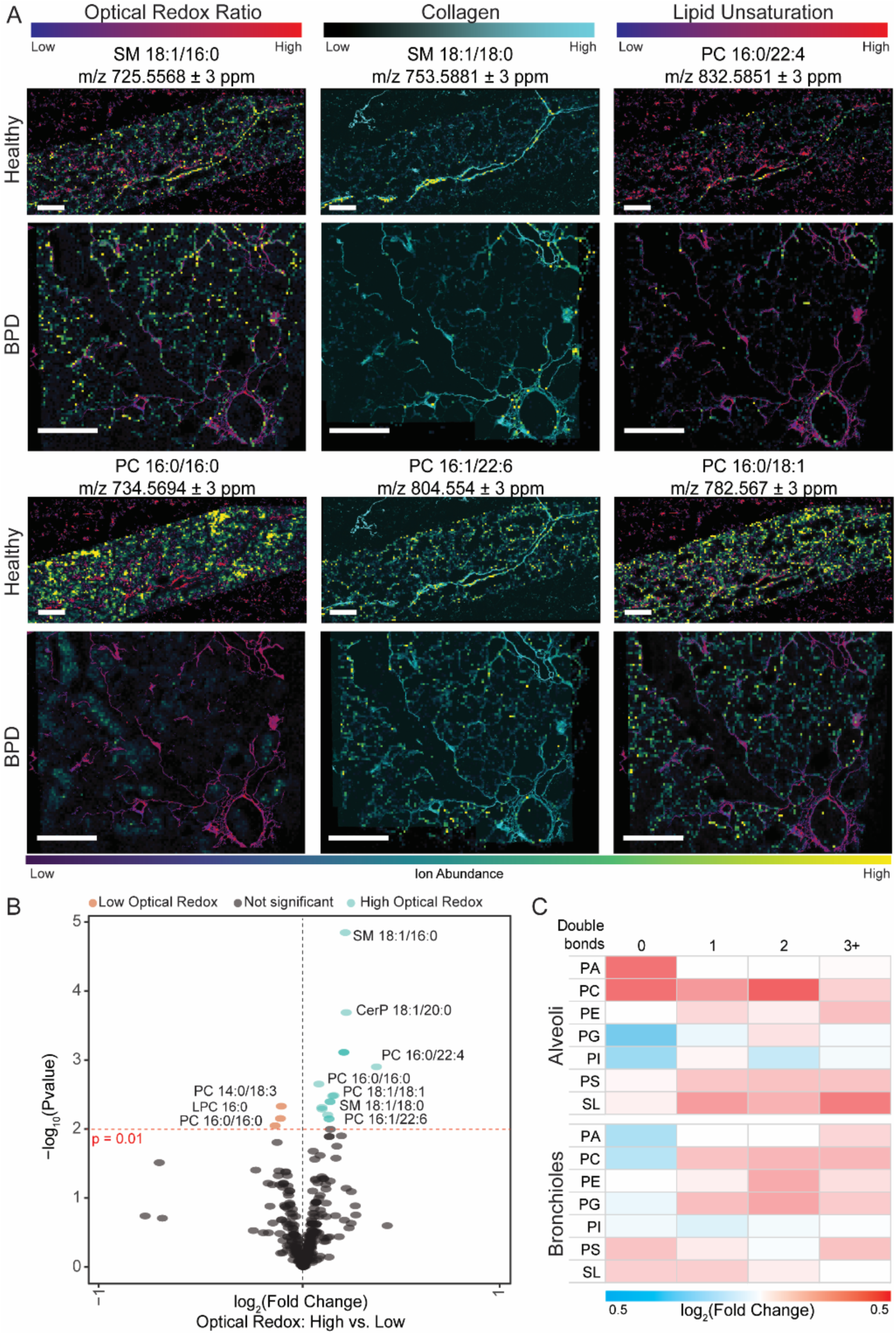
Correlated analysis of co-registered MALDI-MSI and U-FLIP. (A) Co-registered multimodal images of the optical redox ratio, collagen, and lipid unsaturation and representative MALDI-MS images of lipid distributions in the same regions. The optical redox ratio is higher in the vasculature of the tissue than the tissue and displays lower redox in the alveoli regions. This corresponded to a similar increase in SM in the vasculature and increased PC 16:0/16:0 in the alveoli regions. The collagen images show the presence of collagen-rich airways and septa in the lung tissue that corresponded with SM 18:1/18:0 and PC 16:0/18:1. The lipid unsaturation signals provide high resolution information concerning lipid saturation and corresponded with high unsaturation near PC 16:0/22:4 and low unsaturation near PC 16:0/18:1. Scale bars are 500 µm. (B) The volcano plot shows the ions that increase in low and high optical redox regions (p ≤ .01). Ten molecules had significantly increased abundance in high redox regions, and three molecules were increased in Low Redox regions. (C) Heatmaps compare the abundance of double bonds in the Alveoli and Bronchioles compared to all other FTUs. The alveoli contain higher abundances of unsaturated PAs and PCs and unsaturated SLs. The bronchioles displayed slight increases in unsaturation in all lipid species.

In addition to the qualitative function of such images, we used the co-registered image modalities to segment MALDI-MSI data and analyze the regions of interest for enriched lipids. As previously stated, the annotated vessels displayed an increased optical redox ratio compared to the bronchioles (**Figure 7A).** We also observed similar changes in the lipid distributions where the vasculature and arterial regions contained an increased abundance of SM lipids. The registered images allowed us to segment high-redox and low-redox regions and evaluate the ion abundances of high-optical redox and low-optical redox regions. These analyses revealed that ten annotated lipids displayed increased abundances in high-optical redox regions, whereas three lipids increased in abundance in low-optical redox regions (p < 0.01). Of the ten molecules upregulated in high-redox regions, five were sphingolipids, including SM 18:1/16:0, CerP 18:1/20:0, and SM 18:1/18:0. Multiple PCs, including PC 16:0/16:0 and PC 18:1/18:1 that are associated with the surfactant, were also identified in the high-redox regions (**Figure 7B**).

In addition to segmentation based on U-FLIP images, we performed correlated lipid analyses on segmented FTUs. Using U-FLIP, we detected a relatively higher degree of unsaturation in the alveoli compared to the bronchioles. We also evaluated the lipid unsaturation identified by MALDI-MSI. As shown in **Figure 7C**, the saturation of each lipid species varied among FTUs. As suggested by the U-FLIP analysis above, lipid unsaturation increased in the alveoli compared to other regions in the tissue. When correlated with the MALDI analysis, we identified two classes of lipids, PCs and SLs, that increased in saturation in the alveoli. These increases may contribute to the detected increase in lipid unsaturation with U-FLIP. In addition, the bronchioles also displayed a modest increase in the unsaturation of phospholipids. Notably, the MALDI-MSI methods employed here were not optimized for all lipid subtypes, and additional lipid species (e.g., cardiolipins, triglycerides, or diglycerides) may contribute to the increase in unsaturation observed in the multimodal microscopy images.

## Discussion

Multimodal imaging provided valuable multi-domain data to evaluate the complex molecular physiology of organs FTUs and their changes resulting from disease. We applied a newly developed workflow combining MALDI-MSI with U-FLIP. Using each technology, we evaluated the molecular composition of the alveoli, bronchi, and vessels in normal lung tissue. In the bronchi, we observed a relatively higher proportion of longer-chain PCs (**Figure 3A**) and more FA saturation (**Figure 7C**). Noteworthily, longer chain phospholipids provide increased membrane rigidity^43–45^. However, due to the limited number of samples analyzed, our histological annotations do not contain enough granularity to identify airway size to provide broader depictions of the tissue. The analysis of a larger number of samples, originating from different topological regions of the organ would be required to gain further granularity and determine if the chain length of PCs correlates with airway size.

The annotated vessels displayed high abundances of long-chain PCs and SLs (**Figure 3A**). Sphingolipids are implicated in many cellular processes, including molecular transport, immune response, and signaling^46^. The prominent SM 18:1/16:0 is abundant in cellular membranes where it provides membrane fluidity and can aid in communication and extracellular transport^47^. The increased presence of FAD to NADH ratio (**Figure 4D**) in the vasculature may reflect the high energy needs of cells in the vessel walls^48^. Within these high optical redox regions, we also observed a high abundance of SM 18:1/16:0, CerP 18:1/20:0, and SM 18:1/18:0 (**Figure 7b**). These SLs are common in cellular membranes and are often implicated in repair and inflammation^49^.

MALDI-MSI detected PC 16:0/16:0 and PC 16:1/16:0 abundantly within the alveolar parenchyma (**Figure 3A**). Lipids, in particular PCs, make up 90% of pulmonary surfactant, and the most abundant lipid component of surfactant is PC 16:0/16:0 (also known as DPPC)^50^. We compared these observations of normal molecular conditions to changes induced by BPD. Within the BPD airways, we observed an increase in PIs and a decrease in the collagen fiber organization. PI regulation may be affected by the downstream regulation of phosphatidylinositol 3-kinase, an enzyme known to be active during fibrogenesis^51^. The role of the airway collagenous basement membrane is to provide airway stability and rigidity^52,53^. Here, we observed diffuse and unstructured collagen organization in the BPD airways likely to alter airway cells/basement membrane interactions and airway rigidity. This might contribute to the previous observation of reduced passive respiratory system compliance in BPD patients^54^.

MALDI-MSI analysis also revealed decreased PCs and PGs found in BPD alveolar parenchyma. We noted decreased abundances of PC 16:0/16:0 and PC 16:0/16:1, which are common in surfactant. Surfactant is produced by alveolar type 2 (AT2) cells to coat most of the alveolar surface and due to its high packing density, PC 16:0/16:0 provides the surface tension reduction needed at the air-liquid interface of the distal lung during the breathing cycle^55^. In BPD, decreased abundance or function of alveolar type 2 (AT2) cells, may lead to changes in the metabolic profile of both the alveolar epithelium and the pulmonary surfactant^1^. Unfortunately, our MALDI image resolution does not provide differentiation between the surfactant layer and alveolar cells. On the other hand, U-FLIP provided the necessary resolution to detect differences between the outermost layer of the alveoli and the inner tissue layer. We observed an increase in PCs and lipid unsaturation in the innermost layer of the BPD alveoli (**Figure 5).** Many factors can influence the observed changes in lipid composition, including a reduction in surfactant production, tissue remodeling resulting in changes in the availability of circulating lipids during BPD, increased reliance on AT2 for de novo lipid synthesis^1,5,42,56,57^, or an increased prevalence of alveolar macrophages with distinct lipid profile^58^. However, one limitation of this finding is the potential diffusion and removal of soluble biofluids, such as mucus and surfactant, during the steps of embedding and solvent washes before U-FLIP.

## Conclusions

We demonstrate the advantages of employing a multimodal imaging workflow to investigate metabolic differences between lung FTUs from healthy and diseased tissue. Using MALDI-MSI, we identified lipids enriched in four major FTU of healthy lung tissue and then compared the relative abundances of these lipids to those found in a BPD lung noting significant changes in the lipidomic composition. Following MALDI-MSI, we employed U-FLIP to explore broad metabolic changes at higher resolution. This analysis revealed increased lipid unsaturation and optical redox in the vasculature of healthy lung tissue and an overall increase in optical redox in BPD lung tissue. To accurately align serial MALDI-MSI and U-FLIP images, we implemented a novel co-registration strategy called HiMReg. This co-registration allowed us to perform qualitative analysis of the images and further segment the data based on the optical redox ratio images. We discovered that multiple SLs were upregulated in high-redox regions, while a few PCs were upregulated in low-redox regions. Overall, our workflow provides a method for analyzing broad metabolic changes at high spatial resolution, and it permitted us to link these metabolic shifts to specific lipidomic alterations within tissue sections. Continued advancements in multimodal workflows will facilitate the mapping of specific FTUs and their disease-related changes with promise to enhance our understanding of pathological processes in the lung and other tissues.

## Materials and Methods

### Lung Tissue Preparation

Lung tissue was procured through the BioRepository for INvestigation of Diseases of the Lung (BRINDL)^59^. Four agarose inflated, CMC embedded fresh-frozen lung tissue blocks were chosen from the left-upper lobe of a donor with bronchopulmonary dysplasia and an age matched control. Serial tissue sections were cut at 20 µm thickness and thaw mounted on two Superfrost™ Plus Microscope Slides (Fisher Scientific) and two ITO-coated glass slides (Delta Technology). Additional tissue sections (200 µm) were collected in a 1.7 ml Sorenson tube for MPLEx sample preparation and lipid extraction and liquid-chromatography mass spectrometry (LC-MS) analysis.

### MALDI-MSI Sample Preparation and Data Acquisition

Tissue sections were pre-analyzed by autofluorescence microscopy using a Ziess Axio Zoom Microscope. Red (545/572), green (488/509), and blue (353/465) autofluorescence channels were collected along with and a brightfield image. ROIs were selected so that at least one airway, muscularized vessel, and alveolar region was analyzed from each tissue section. A M5-Sprayer (HTX Technologies) was used for matrix application, where a N_2_ pressure of 10 psi, track spacing of 3 mm, and a 40 mm distance between the nozzle and sample was maintained for preparation of all samples. For positive ion mode analysis, 2,5-dihydrobenzoic acid (DHB; 40 mg/ml in 70% MeOH:H_2_O) was sprayed at a rate of 50 µl/min and 75 °C for 12 passes. For negative ion mode, N-(1-Naphthyl)ethylenediamine dihydrochloride (NEDC; 7 mg/ml in 70% MeOH:H_2_O) was applied at 120 µl/min and 75 °C for 8 passes.

MALDI-MSI analysis was performed using a Bruker 12 tesla Fourier transform ion cyclotron resonance mass spectrometer (FTICR-MS) in positive and negative ion mode. Both modes were tuned using sodium trifluoroacetate (1 µg/ml) and maximal transmission and detections were optimized based on signals in the 600–900 m/z range. The positive ion mode method used 120 laser shots at 2000 Hz and had a mass resolving power ∼180,000 at 400 m/z. In negative ion mode 100 laser shots were applied at 2000 Hz and ∼240,000 at 400 m/z mass resolving power was achieved. When performing MALDI-MSI on glass slides, the slides were mounted using cooper tape on a Bruker target plate.

### MALDI-MSI Data Processing

MALDI-FTICR-MSI data was imported into Bruker SCiLS Software (Version 2024b) and converted into the imzML format. The resulting imzML and ibd files were uploaded to METASPACE^60^ for automated molecular annotation against the SwissLipids, LipidMaps, and internal LC-MS/MS lung databases. Lipid identifications from Swisslipids (FDR ≤ 20%) and the custom LC-MS/MS database were downloaded and imported back into SCiLS for data integration and visualization. Of the 280 lipids putatively annotated by the Swisslipids database (FDR ≤ 20%) and confirmed using annotations against an internal LC-MS/MS database, 108 species were present in over 70% of the analyzed regions. These 280 molecular annotations were imported into SCiLS; and histology images and UFLIP images were manually aligned with the MALDI images within SCiLS for visualization and analysis.

MALDI-MSI, histology segmentations, and UFLIP image segmentations were exported through SCiLS API (2024b) and analyzed in ‘RomicsProcessor’ (V1.6, https://github.com/PNNL-Comp-Mass-Spec/RomicsProcessor), an open-source R package for mass spectrometry data analysis. Ion intensities were median-centered and a log_2_ transformation was applied. A two-tailed Student’s t-test was performed to compare each annotated region (i.e., BPD alveoli, healthy alveoli, BPD bronchi, healthy bronchi, etc.) to all other regions. Log_2_ fold intensity changes between regions, and p-values were computed for each.

### Bulk Lipidomics

An internal lipid database was created using LC-MS/MS data collected from 200 µm serial tissue sections. Briefly, on each section the metabolite, protein, and lipid extraction (MPLEx) protocol was performed, as described previously^61–63^. Additional details can be found in the supplemental information. Confident lipid identifications were made using the in-house developed identification software LIQUID, where the tandem mass spectra were examined for diagnostic ion fragments along with associated hydrocarbon chain fragment information^64^.

### U-FLIP

An upright laser-scanning microscope (DIY multiphoton, Olympus) with a 25× water objective (XLPLN, WMP2, 1.05 NA, Olympus) was applied for near-IR throughput. Synchronized pulsed pump beam (tunable 720–990 nm wavelength, 5–6 ps pulse width, and 80 MHz repetition rate) and Stokes (wavelength at 1032nm, 6 ps pulse width, and 80MHz repetition rate) were supplied by a picoEmerald system (Applied Physics & Electronics) and coupled into the microscope. The pump and Stokes beams were collected in transmission configuration by a high NA oil condenser (1.4 NA). A high O.D. shortpass filter (950 nm, Thorlabs) was used that would completely block the Stokes beam and transmit the pump beam only onto a Si photodiode for detecting the stimulated Raman loss signal. The output current from the photodiode was terminated, filtered, and demodulated in X with a zero-phase shift by a lock-in amplifier (HF2LI, Zurich Instruments) at 20 MHz. The demodulated signal was fed into the FV3000 software module FV-OSR (Olympus) to form the image during laser scanning. All SRS images were obtained with a pixel dwell time 20 µs and a time constant of 15 µs. Laser power incident on the sample is approximately 40 mW. SHG was used to capture Collagen types 1-3 images. The 1031 nm stokes laser described above, with 300 mW and a dwell time of 8 µs per pixel, was used with 3-frame averaging. Backscattered SHG signals were filtered using 465 nm filter. NADH and Flavin autofluorescence images were captured using the 800 nm pump beam with 350 mW and a pixel dwell time of 10 µs/px with 3-frame averaging. Backscattered signals were filtered using a dual filter cube of 460 nm and 515 nm. All wide-view tile-stitching was controlled by the Fluoview software (Olympus). Hyperspectral SRS (hSRS) images for chemical composition analyses were acquired with 60-frame image stack, and other parameters are the same as single SRS image.

### Histology Imaging

The same section imaged with MALDI-MSI and U-FLIP when mounted on glass slides were finally stained by a standard hematoxylin and eosin (H&E) staining protocol. Each slide was rinsed with 70% EtOH and H_2_O for 1 min each. Then the slides were stained for 30 s in Mayer’s hematoxylin solution (1 g/L; Sigma-Aldrich). Slides were rinsed with H_2_O and dipped in a bluing solution (American Master Tech Scientific) for 20 s, followed by incubation in H_2_O, 70% EtOH, and 95% EtOH for 30 s each. Lastly, slides were dipped in eosin (1%, Sigma-Aldrich) 4× and dipped in 95% EtOH twice, 100% EtOH twice, and xylene for dehydration. Each tissue section was cover slipped and imaged using an Aperio ScanScope XT at 20× magnification. Images were imported into QuPath (v0.3.8)^65^, where the alveoli, bronchi, bronchioles, and vessels were manually segmented.

### Hierarchical Multimodal Registration Network (HiMReg)

HiMReg implements a two-stage registration optimization combining hierarchical affine and diffeomorphic transformations. The network leverages multi-scale from coarse-to-fine processing and GPU acceleration through PyTorch to achieve efficient and accurate alignment of multimodal images.

Stage 1: Hierarchical Affine Registration. The affine transformation 𝑇_𝐴_ maps source coordinates x = (𝑥, 𝑦) to target coordinates x^′^ through:

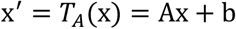

where A is a 2 × 2 matrix encoding rotation and scaling, and b is a translation vector. The optimization minimizes:

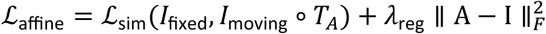

where ℒ_sim_ is a similarity metric (mutual information or local normalized cross correlation), 𝜆_reg_ is a regularization parameter, and ∥∥⋅∥∥_F_ denotes the Frobenius norm.

Stage 2: Hierarchical Diffeomorphic Registration. The diffeomorphic stage refines local deformations through a displacement field v(x). The transformation 𝜙 maps the spatial coordinates directly:

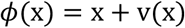

where 𝜙(x) represents the spatial transformation mapping initial coordinates x to their deformed positions, and v(x) is the displacement field that defines the deformation. To ensure the transformation is diffeomorphic (smooth and invertible), we regularized the displacement field gradient and the final optimization objective combines similarity and regularization terms

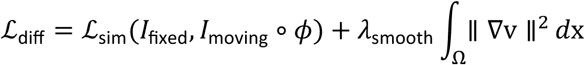

where the first term measures the similarity between the fixed image and the transformed moving image, while the second term enforces smoothness of the displacement field through its spatial gradient. The smoothness weight 𝜆_smooth_ controls the trade-off between image alignment accuracy and transformation regularity to avoid over transformation.

Each registration stage employs a hierarchical approach with 𝐿 resolution levels. At each level 𝑙, images are downsampled. The optimization begins at the coarsest level (highest downsampling factor) and progressively refines the transformation parameters 𝜃_𝑙_ as the resolution increases:

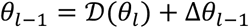

where 𝒟 adapts the transformation parameters from a coarser level 𝑙 to a finer level 𝑙 − 1, and Δ𝜃_𝑙–1_represents the refinement at the finer scale. This hierarchical strategy first captures large-scale deformations at coarse resolutions (e.g., 8× downsampling) before refining local details at finer scales (e.g., 4×, 2×), ultimately reaching the original resolution.

Due to the multimodal nature of our registration task. We employed mutual information (MI) as the primary similarity metric ℒ_sim_. MI is particularly suitable for multimodal registration as it makes no assumptions about the intensity relationships between different imaging modalities. The MI between two images X and Y is defined as:

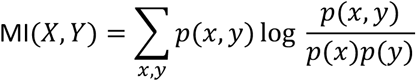

where 𝑝(𝑥, 𝑦) is the joint probability distribution of the intensities in images X and Y, and 𝑝(𝑥) and 𝑝(𝑦) are their respective marginal distributions. MI measures the statistical dependency between the intensities of the two images, reaching its maximum when the images are optimally aligned.

The registration evaluation was also assessed using the Dice similarity coefficient (DSC), which measures the spatial overlap between the transformed moving image and the fixed image:

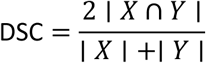

where ∣ 𝑋 ∣ and ∣ 𝑌 ∣ represent the cardinalities of the two segmented regions, and ∣ 𝑋 ∩ 𝑌 ∣ is the size of their intersection. The DSC ranges from 0 to 1, providing a quantitative measure of registration accuracy.

### U-FLIP Analysis

All image analysis was performed using custom Python scripts. For TPF data, NADH and FAD images were first corrected for background using a threshold-based mask. The normalized optical redox ratio was calculated pixelwise as FAD/(NADH + FAD). Regions with signal intensity below the noise threshold were excluded from the analysis. SHG images were analyzed to quantify collagen fiber organization using a custom pipeline. Images were pre-processed using Gaussian filtering (σ=1) to reduce noise while preserving fiber structures. Binary masks were generated using Otsu thresholding, followed by morphological operations to remove small artifacts. Fiber density was calculated as the ratio of collagen-positive pixels to total tissue area. Local fiber thickness was measured using distance transform methods on the skeletonized fiber network. Hyperspectral SRS data were processed using a multi-step workflow. Raw spectral data underwent baseline correction to remove water background. Spectra were then normalized using maximum intensity normalization and smoothed with a Whittaker filter (λ=0.2). K-means clustering (k=3) was applied to the processed spectra to identify chemically distinct regions. The lipid unsaturation ratio was calculated from intensity-normalized SRS images as I_3011_ / (I_2880_+ I_3011_). For spatial analysis of FTUs, ROIs were manually segmented using QuPath software. Then 20,000 pixels from each FTUs were selected randomly for the pixel-wise quantitative analysis.

## Code Availability

Demo data and source code for HiMReg are available at https://github.com/Zhi-Li-SRS/HiMReg.

## Data Availability

All MALDI-MSI images are located on METASPACE https://metaspace2020.org/project/gorman-2025

## Acknowledgements.

This work was supported by the National Heart, Lung, and Blood Institute (NHLBI) Molecular Atlas of Lung Development Program Human Tissue Core (LungMAP HTC) and LungMAP BioRepository for INvestigation of Diseases of the Lung (BRINDL) through grants U01HL122700 and U01HL148861 (to GSP), and U01HL148860 (G. Clair) and by the NIH Common Fund grant U54 HL165443-01 (to GSP, with Co-Investigators GC, and CA). Part of this work was performed in the Environmental Molecular Science Laboratory, a U.S. Department of Energy (DOE) national scientific user facility at Pacific Northwest National Laboratory (PNNL). Battelle operates PNNL for the DOE under contract DE-AC05-76RLO01830. The opinions expressed in this article are the authors’ own and do not reflect the view of the NIH, the Department of Health and Human Services, or the U.S. government. We are very grateful for the generosity of the donor families and honor their loss.

