## Supplemental Data for "A multimodal imaging approach for imaging the metabolic changes resulting from bronchopulmonary dysplasia"

*Corresponding to:

Lingyan Shi,

Christopher Anderton,

**Supporting Information**

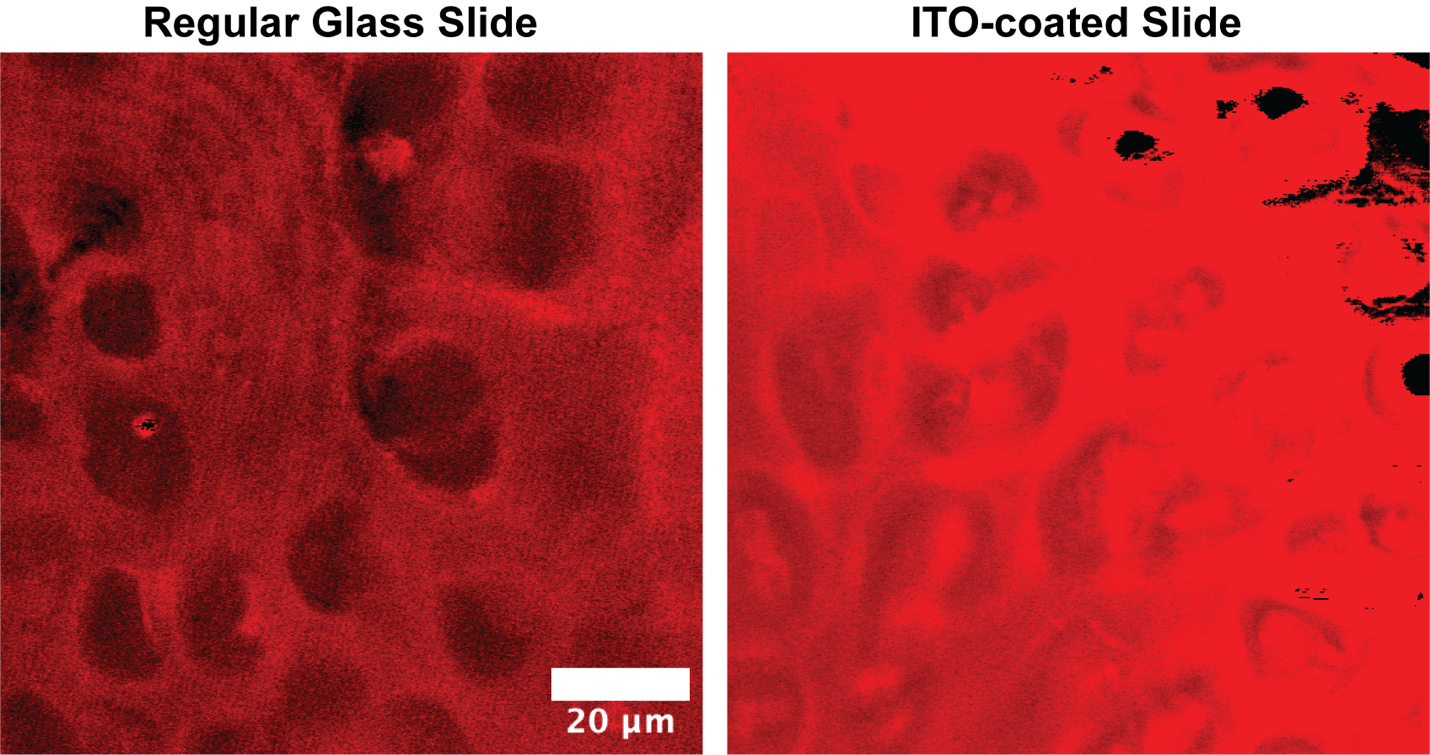

**Figure S1.** **Comparison of SRS protein channel images acquired from human lung sample on regular glass and ITO-coated slide**. SRS imaging of the protein channel demonstrates significant differences in image quality between standard glass and ITO-coated slide. Images acquired on ITO-coated slide exhibit consistent oversaturation and substantially reduced signal-to-noise ratio across all Raman shifts.

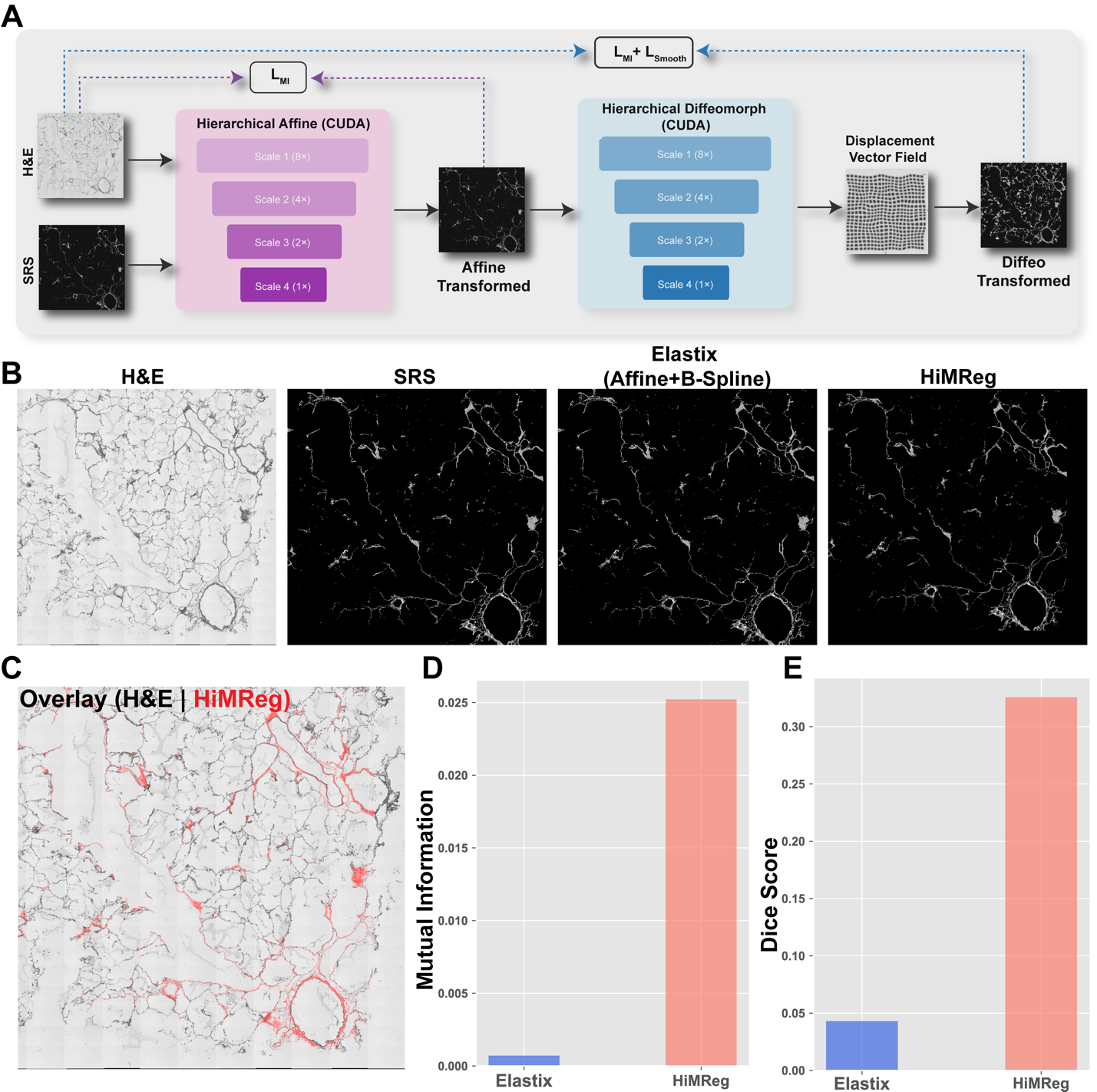

**Figure S2. Hierarchical Multimodal Registration network (HiMReg) architecture and performance evaluation.** **(A)** Schematic overview of the HiMReg architecture showing the two-stage registration process: hierarchical affine registration (purple) followed by hierarchical diffeomorphic registration (blue). Each stage employs a customized pyramidal approach with GPU acceleration. **(B)** Visual comparison of registration results between original H&E and SRS images using Elastix (Affine+B-Spline) and HiMReg methods. **(C)** Overlay visualization of H&E (grayscale) and HiMReg-registered SRS (red) images demonstrating precise alignment of tissue structures. **(D,E)** Performance comparison between Elastix and HiMReg using mutual information (D) and dice (E) metrics, showing superior registration accuracy achieved by HiMReg.

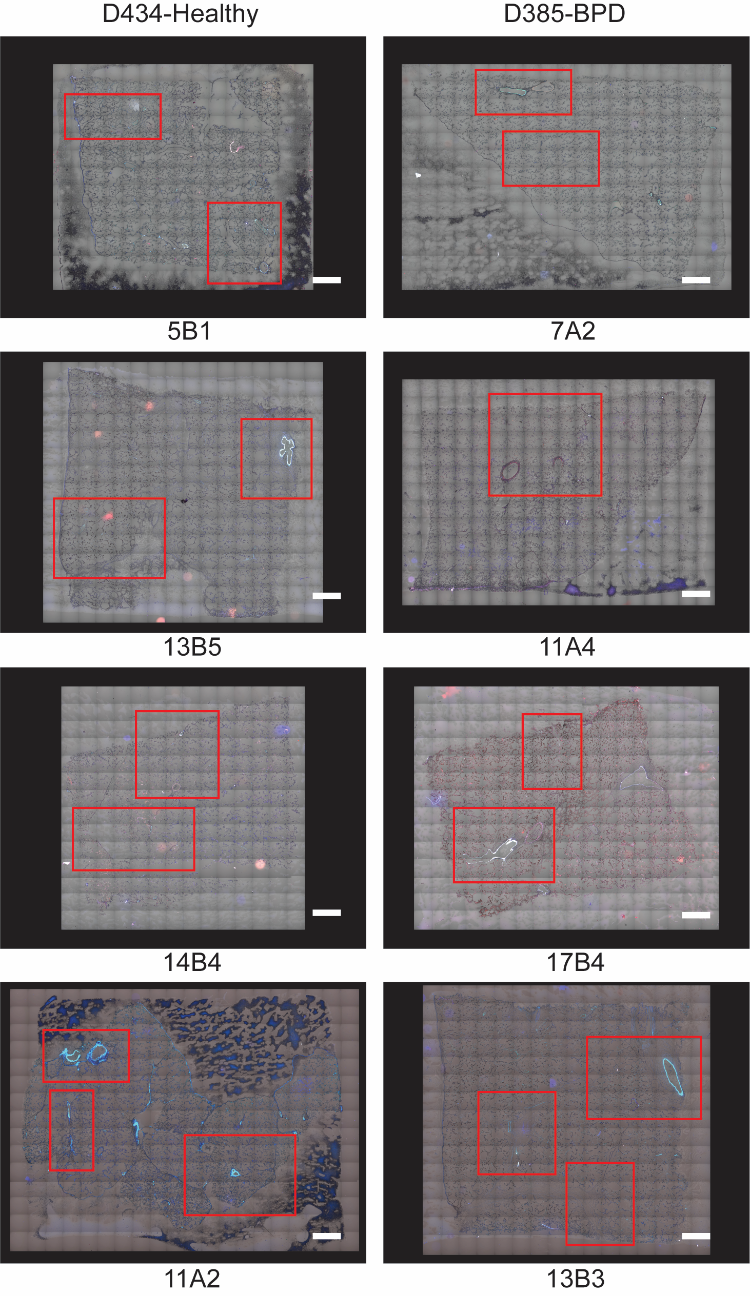
**Figure S3. Autofluorescence images of each tissue analyzed section.** Autofluorescence images showing the red, green, blue, and transmitted light. ROIs were selected for MALDI-MSI analysis at 35 µm containing bronchi, bronchioles, vessels, and alveolar parenchyma (red boxes).

**Tabel S1.** All human lung tissue block IDs

| **Donor ID** | **Block ID** | **Condition** | **Cause of Death** | **GA at Birth (weeks)** | **Postnatal Age at Demise (months)** | **Histopathology** |
| --- | --- | --- | --- | --- | --- | --- |
| D385 | LUL-13B3  LUL-17B4  LUL-13B5  LUL-7A2 | Broncho-pulmonary Dysplasia | Broncho-pulmonary Dysplasia | 25 | 7 | Chronic lung disease of prematurity, reduced alveolarization, enlarged simplified airspaces, patchy early interstitial fibrosis and extension of smooth muscle into lobules, Focal bronchial squamous metaplasia |
| D434 | LUL-11A2  LUL-11A4  LUL-14B4  LUL-5B1 | No known lung disease | Cardiovascular/ natural causes | 40 | 2 | Normal lung structure and alveolar growth. Patchy minimal acute and chronic airway inflammation |
